# Host environmental conditions induce small fungal cell size and alter population heterogeneity in *Cryptococcus neoformans*

**DOI:** 10.1101/2020.01.03.894709

**Authors:** Xin Zhou, Hanna Zafar, Poppy Sephton-Clark, Sally H. Mohamed, Ambre Chapuis, Maria Makarova, Donna M. MacCallum, Rebecca A. Drummond, Ivy M. Dambuza, Elizabeth R. Ballou

## Abstract

Fungal morphology significantly impacts the host response. Filamentation and tissue penetration by *Candida* and *Aspergillus* species are essential for virulence, while growth as a yeast allows the thermal dimorphic fungi *Coccidiodes, Histoplasma,* and *Talaromyces* to reside inside phagocytes and disseminate. The basidiomycete *Cryptococcus neoformans* exhibits an unusual yeast-to-titan transition thought to enhance pathogenicity by increasing fungal survival in the host lung and dissemination to the central nervous system. In a common laboratory strain (H99), *in vitro* and *in vivo* titan induction yields a heterogenous population including >10 μm titan cells, 5-7 μm yeast cells and 2-4 μm titanides. Previous reports have shown that titan cells are associated with enhanced virulence and the generation of aneuploid cells that facilitate stress adaptation and drug resistance, while small (>10 μm) cells are associated with increased dissemination. However, the relationship between titan cells, small cells, and titanides remains unclear. Here, we characterize titanides and small cells in H99 and three clinical isolates and show that titanides share the lipid membrane order of their titan mothers and the G_0_ quiescent-like DNA staining of mating spores. In addition, we show that both titanizing and non-titanizing isolates exhibit altered capsule structure and PAMP exposure over time during *in vitro* culture, and generate aneuploidy *in vivo*.

**Author summary:** The human fungal pathogen *Cryptococcus neoformans* causes 200,000 HIV-associated deaths each year. In the lung, *Cryptococcus* makes an unusual yeast-to-titan morphological switch that contributes to disease development by altering immune polarization and introducing aneuploidy underlying host stress and drug resistance. Specifically, a proportion of 5 um haploid yeast endoreduplicate and swell, converting to large (> 10 um) polyploid titan cells that can then produce genetically distinct daughter cells. We recently developed an *in vitro* protocol for inducing large titan cells and additionally observed a novel small “titanide” cell type. Here we investigate the nature and origin of these small cells, demonstrating that they emerge during both *in vitro* and *in vivo* mouse-passaged titan induction in the well characterised lab strain H99 and are also apparent in a titanizing clinical isolate, Zc8. We show that these titanide cells share features with titan mothers (lipid order) and with spores produced during heterothalic mating. Finally, we show that the capacity of clinical isolates to produce both titan and titanide cells impacts aneuploidy and the emergence of drug resistance *in vivo*.

## Introduction

*Cryptococcus neoformans* is a human fungal pathogen causing life-threatening pneumonia and meningitis in immunocompromised and immunocompetent individuals [1–3]. In the host lung, inhaled *C. neoformans* spores and yeast proliferate and can undergo an unusual morphogenic switch from 5-7μm haploid yeast to large (>10 μm) polyploid titan cells with altered cell wall and capsule organization [4–7]. This is a form of fungal gigantism, with individual cells reaching 50 to 100 μm [4–6, 8–10]. Titan cells also produce aneuploid small (<10 μm) daughter cells, although the importance of these in disease progression is unclear [3, 11–14]. Titan cells enhance the dissemination and virulence of *C. neoformans* in animal and mini-host models of infection [4, 5, 15–18]. Yet, titan cells block phagocytosis, and phagocyte-mediated dissemination, of non-titan cells. Mutants deficient in titan cells exhibit reduced virulence, and mutants that produce fewer small cells disseminate more slowly [15, 19–21].

A pressing clinical question is how titan cells influence patient outcome in a clinical setting. Clinical isolates vary widely in their expression of known virulence factors including capsule, melanin, and urease, a result of the wide range of infecting genotypes even in relatively localized settings and within individual patients [13, 22–26]. Given this diverse virulence portfolio, it is unsurprising that a screen of clinical isolates demonstrated that fungal cell size alone is insufficient to explain *in vivo* outcomes [22]. However, two recent analyses of clinical isolates highlighted a role for small cells in disease progression. Following long term growth in capsule-inducing medium, Fernandes *et al*. observed either giant (>15 μm including capsule) or thick-walled micro (<1 μm) cells in *C. tetragattii* and *C. neoformans* isolates, respectively, and correlated this with CD4 clinical outcomes [13]. Rare *C. neoformans* isolates that produced both giant and micro cells were highly virulent. Mukaremera *et al*. also observed an inverse correlation between virulence in humans and the presence of large cells following culture in DMEM+serum, conditions in which the virulent, titanizing isolate KN99α failed form titan cells [22]. Denham et al showed the emergence of small cells in the lung over the course of infection and showed that small cells predominate in the brain [19]. Together both studies suggest a role for small cells in infection progression, however the link between small and titan cells remains undetermined. By studying H99 titan-induced cultures, we identified a distinct small cell type which we termed titanides [14]. Titanides have distinct morphological properties: oval cell shapes, small cell size (2-3 μm) and thin cell walls [14]. These are morphologically distinct from previously described micro cells (round, thick cell wall, <1 μm) and drop cells (5μm), both of which have been associated with *in vivo* infection [8, 27, 28].

Genetic and genome-wide analyses in *C. neoformans* H99 have shown that the yeast-to-titan transition is a complex phenotype regulated by multiple interacting pathways[14, 20, 29–31]. Titan cells have been observed in *C. neoformans* VNI, VNII, VNB, *C. deneoformans* and *C. gattii* clinical and environmental isolates during growth in established *in vitro* titan-inducing conditions that recapitulate the host environment [14, 29, 31, 32]. However, in these conditions, we and others also observed a striking variation in size range and morphology regardless of the presence of classic titan cells (>10 μm cell body) [14, 29, 31]. Given evidence of induced population size heterogeneity across and within species, as well as evidence demonstrating conserved genetic regulation of titanization across species, we hypothesized that even in isolates that do not exhibit classic >10 μm “titan” cells, exposure to *in vitro* titan-inducing conditions might trigger important changes in titan-related phenotypes (size distribution, cell wall organization, ploidy, capsule), with implications for fungal virulence.

To investigate this, we examined the impact of *in vitro* titan-inducing conditions on four representative clinical isolates: H99, Zc1, Zc8, and Zc12, selected based on their genetic relatedness and their range of titan phenotypes: titanizing (H99, Zc8) or non-titanizing (Zc1, Zc12). Here, we fully characterized these isolates and find that, like titanizing strains, ‘non-titanizing’ strains also display population heterogeneity in size distribution, cell wall organization, and DNA content following exposure to inducing conditions. We identify titanide cells both during *in vitro* culture and *in vivo* and demonstrate that titanides are the progeny of polyploid titan cells, sharing their lipid membrane organization. Moreover, we show that titanides share features of quiescent cells and spores. In addition, we find that the ‘non-titanizing’ Zc12 isolate, despite being defective in producing titan and titanide cells, do produce small cells and can generate aneuploid offspring *in vivo*. Finally, we observe that *in vivo* aneuploidy does not directly correlate with drug resistance. Together, this work confirms the importance of small cells for disease progression, identifies titanides as a specific subset of small cells originating from titan mothers, and raises questions about the role of population heterogeneity in disease progression.

## Results

In this study, we examine clinical isolates previously collected from the ACTA Lusaka Trial in Lusaka, Zambia [26, 33]. We previously showed that Zambian clinical and environment strains collected by Vanhove *et al*. vary widely in their capacity to form titan cells, irrespective of originating niche or genotype [14]. Here, we further characterize titanization in four strains-H99, Zc1, Zc8 and Zc12-selected based on their capacity to form titan cells *in vitro*. *In vivo* induced titan cells are characterized by four defining features: enlarged cell sizes, high ploidy, tightly compacted capsule, and the presence of a single, large vacuole [6, 34]. Using this definition, established by Nielsen and Zaragoza, we previously classified Zc8 as titanizing, and Zc1 and Zc12 as non-titanizing [6, 14]. We also showed that Zc1 exhibits reduced virulence compared to H99, and that all four isolates produce melanin to comparable levels [14].

### *In vitro* titan inducing conditions trigger a range of cell size and DNA content changes across four clinical isolates

All four strains are of clinical origin, have been fully sequenced and genotyped, and are genetically closely related (VNI). During standard *in vitro* YPD culture, cells were uniform in size, generally round with no defects in budding, and cell size was similar across and within all strains. To more fully characterize the impact of titan induction on all four strains, we first examined cell size distribution across the four clinical isolates under *in vitro* titanizing conditions. Consistent with our previous classification, H99 and Zc8 produce highly heterogeneous populations comprising titans (>10 μm), yeasts (>4μm and <10μm), and titanides (>1 μm and <3 μm oval cells), while Zc1 and Zc12 populations are more uniform (Fig 1A). After 24 hours, cells >10 μm account for about 15% of the H99 total population, but only approximately 2% in Zc8 (Fig 1B). Moreover, induced non-titan Zc8 cells (<10 μm) are smaller on average than induced non-titan H99 cells (3.80 μm ± 0.10 vs 7.15 μm ± 0.14 SEM, p<0.0001, n>200), and are smaller on average than Zc1 or Zc12 induced cells (4.44 μm ± 0.05, 4.57 μm ± 0.06 SEM, respectively, p<0.0001, n>200).

**Fig 1.**
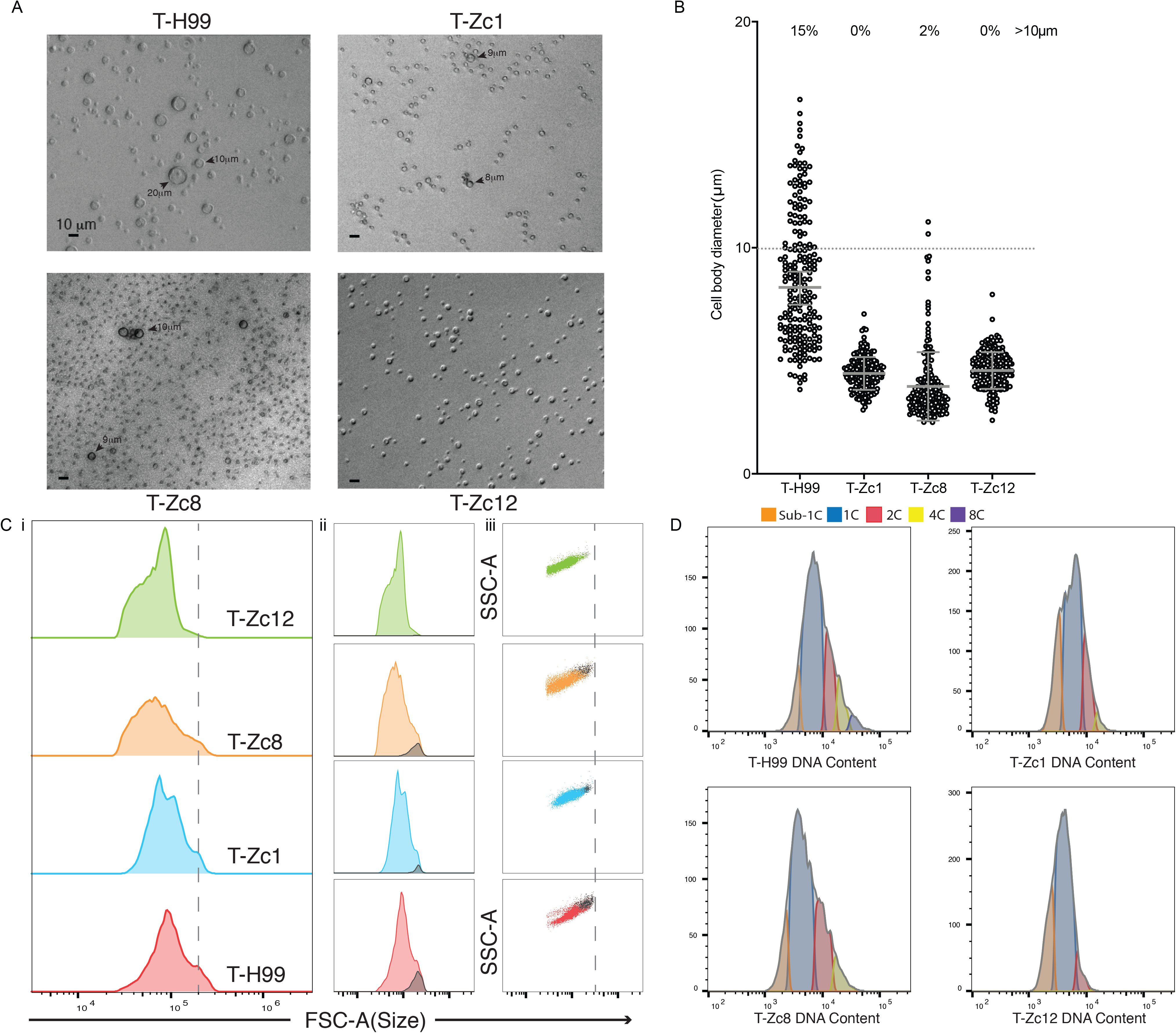
Titan cells formation varies among clinical isolates *in vitro*. (A) *C. neoformans* cells from all indicated strains after 24hrs titan induction, Scale bar: 10μm. (B) Cell body sizes of titan induced cells (n>200). (C) Flow cytometry for cells after titanization with FSC-A as a proxy for cell size. i) Histograms for cell size (FSC-A) are shown for all indicated isolates with a dotted line indicating the distinct shoulder of size increase. ii) The gray sub-population indicates the size threshold identified for cells with DNA content >4C. iii) FSC-A vs SSC-A scatter plots for all indicated isolates with a dotted line indicating the threshold for H99 cell sizes (D) Flow cytometry analysis for DNA content after staining with DAPI. 1C and 2C populations were identified relative to YPD grown yeast cells from matched parent strains (Fig S1A).4C peaks were established according to doubled MFI of DAPI staining relative to 2C. “T” indicates pre-titanized cells. For all figures, data for a representative biological replicate is presented (n>3).

Given the low frequency of titans detected in induced Zc8 cultures, which can easily be missed when relying on cell counts with small numbers (n=200), we assessed cell size spread and DNA content across the population of a large number of cells by flow cytometry (up to n=10,000). Using this approach, for H99 we observed population structure characteristic of typical and titan populations, with a minority sub-population of cells distinguishable as a shift to the right by forward scatter (FSC, a common proxy for cell size) (Fig 1Ci, n>8000) and DNA content >4C (12.3%, Fig 1D, see Fig S1A for matched YPD controls). However, while 6.4% of induced Zc8 cells had DNA content >4C, this FSC shift was not apparent as H99 (Fig 1Cii). In addition, the median FSC of cells with >4C DNA content was lower than in the H99 population, suggesting these Zc8 polyploid cells may be smaller than H99 polyploid cells. Despite this overall reduction in frequency, a similar stepwise population of polyploid cells with a long tail to the right was apparent for Zc8 (Fig 1D). Unlike H99, the vast majority of cells in titan-induced Zc1 and Zc12 populations had DAPI staining consistent with 1C DNA content. However, when we closely examined DNA content, we did identify rare polyploid 4C cells in Zc1 isolates (2.8%) and Zc12 (<1%). These cells were similar in FSC to H99 4C cells, suggesting the presence of rare titan cells in the Zc1 and Zc12 cultures, which likely require acquisition of larger sample size in order to detect (Fig 1C iii). These data suggest that exposure to titan inducing conditions has a general impact on both clinical isolate ploidy and cell size.

### Induced cells exhibit changes in PAMPs exposure

*C. neoformans* titan and typical cells recovered from mouse lung have different cell wall composition, with an increase in chitin and mannose (including mannan, mannoproteins, and capsular mannans) observed in cells >10 μm [7, 18]. We therefore asked whether similar differences could be observed in *in vitro* induced cells. Chitin (Calcofluor white, CFW), chitosan (EosinY), and cell wall mannan (ConA) were measured following exposure to inducing conditions for 24 hours. Microscopy and flow cytometry revealed an overall decrease in PAMP exposure following titan induction relative to YNB (Fig 2A, 2Bi, H99 representative plot). Gating on positively staining cells only, chitin, mannan, and chitosan staining correlated with cell size, with larger cells staining more intensely than smaller cells, the majority of which were below the threshold of detection (Fig 2Bii). Moreover, distinct high chitin populations were apparent in H99, Zc1, and Zc8 cultures (Fig S2). Quantitative analysis by flow cytometry using the same approach to identify titan cells as taken in Fig 1C again revealed a sub-population of large cells in all four induced cultures (Fig S2). Similar to *in vivo*-derived H99 titan cells, *in vitro*-induced H99, Zc1, Zc8, and Zc12 cells displayed a distinct CFW-high population that correlated with the largest cells (Fig 2B, Fig S2) [7, 18]. The largest cells in all isolates had comparable chitosan and cell wall mannan staining to typical cells (Fig S2). Moreover, in long-term induced cultures (72 h), overall size heterogeneity was more apparent, with increased numbers of small cells and distinct high and low chitin peaks present in both Zc1 and Zc12 populations (Fig 2C). Together, these data suggest that changes in cell wall composition occur during low density growth in serum regardless of induction of cells larger than 10um.

**Fig 2.**
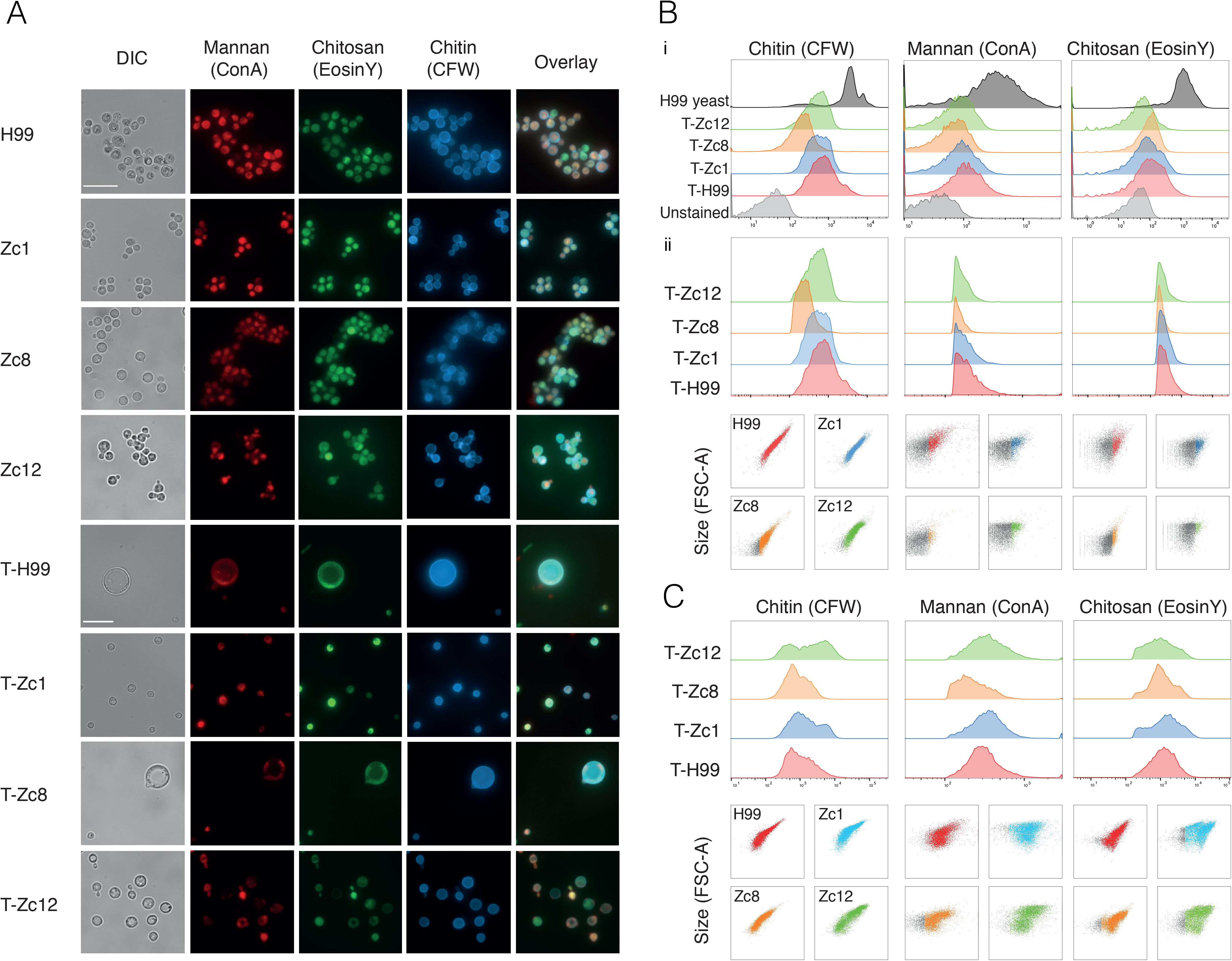
Non-titanising strains also exhibit population heterogeneity. (A) PAMPs staining for all indicated strains cells from both YNB yeast and 24hrs titan induction, Mannan (ConA, Red), Chitosan (EosinY, Green), Chitin (CFW, Blue). YNB grown yeast scale bar 5μm; titanized cells scale bar 10 μm. (B) Stained cells from (A) were analysed by flow cytometry to determine MFI indicating exposure of Chitin (CFW, right); Mannan (ConA, center); and Chitosan (EosinY, left). (C) Flow cytometry analysis of PAMPs staining for indicated cells from long-term(72hrs) titan inductions with MFI indicating exposure of Chitin (CFW, right); Mannan (ConA, center); and Chitosan (EosinY, left). “T” indicates pre-titanized cells.

### Growth in titan inducing conditions can alter virulence relative to yeast-phase growth

Next, we assessed relative virulence of the four isolates using a wax moth larva model [35, 36]. When pre-grown in YPD, the non-titanizing isolate Zc12 was significantly more virulent than the other three isolates (n=10, Gehan-Breslow-Wilcoxon test, p=0.00070) (Fig 3A), killing a majority of larva within 5 days, whereas a majority of Zc8 infected larvae survived to pupation [37]. However, when cells were pre-grown for 24 hr in titan-inducing conditions prior to infection, H99, Zc1, and Zc8 became as virulent as Zc12 (Fig 3A) [14]. Comparing the impact of inducing condition on the change in virulence revealed that no change was observed for Zc12, but induced Zc1, Zc8, and H99 cells increased in virulence relative to un-induced cells (Fig 3B). The observation that exposure to titan inducing conditions does not increase the virulence of Zc12 raised the question of what impact titan inducing conditions might have on Zc1 to alter its virulence.

**Fig 3.**
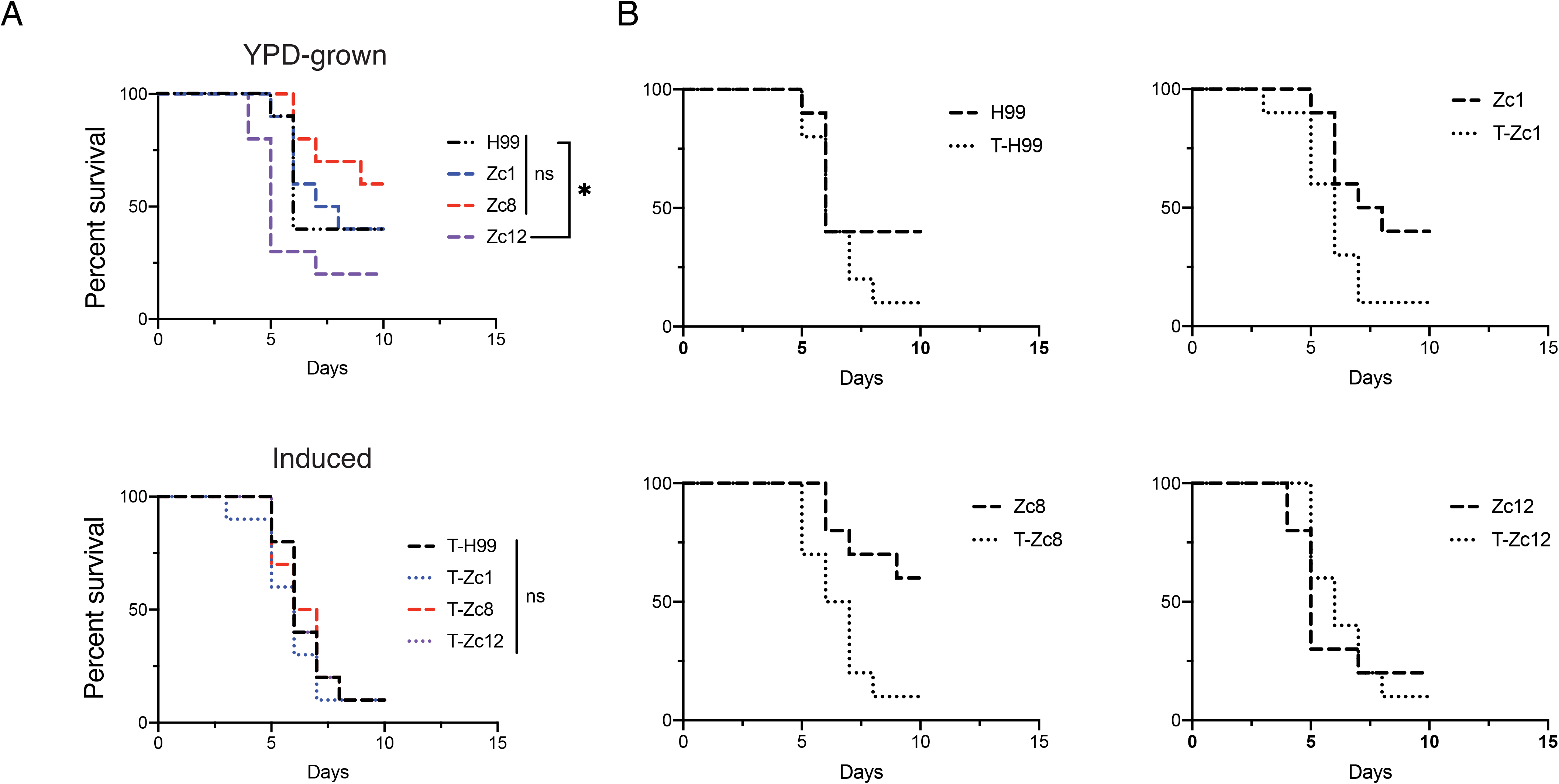
Disease outcomes varied from basal to titan-induced conditions on worm model. (A)Survival curves of worms infected by YPD-grown or titan induced cells of four indicated strains. (B) Comparison of survival curves of worms infected with cells from both conditions. “T” indicates pre-titanized cells. For all figures, data for a representative biological replicate is presented (n>3).

### Cell density is sufficient to change capsule size and structure during capsule induction

To further explore the differences in disease outcome resulting from titan induction, we examined capsule structure from yeast-phase and titan-induced stages. The *C. neoformans* cell wall is covered by a protective capsule, which not only impairs phagocytosis by macrophages, but also masks PAMPs from being engaged by host PRRs[38]. We first examined growth during incubation in 10% FCS at high cell density (OD_600_ 0.1), conditions established to induce robust capsule but not titan cells [14]. Capsule was visualized using india ink and the capsule to cell body ratio measured (Fig 4B, C). There was no relationship between wax moth virulence and capsule size: H99 and Zc12 showed an overall larger capsule size compared to Zc1 and Zc8 (Fig 4B) (p<0.0001, n>100). We previously showed that the anti-capsule IgM antibody Crp127 binds O-mannosylation sites in capsular polysaccharide. In standard capsule inducing conditions, Crp127 binding was highly heterogeneous across and within all four strains, with both stained and unstained cells apparent (Fig 4C), consistent with our previous reports assessing binding during YPD and DMEM-induced growth [39]. As assessed by flow cytometry, H99 spanned unstained to high binding, with 51.8% of cells showing intermediate or high binding. For Zc8, the vast majority stained with intermediate Median Fluorescence Intensity (MFI (67.7%). For Zc1 the majority of cells are unstained (86.4%), while 48.1% of cells from Zc12 exhibited intermediate binding (Fig 4C).

**Fig 4.**
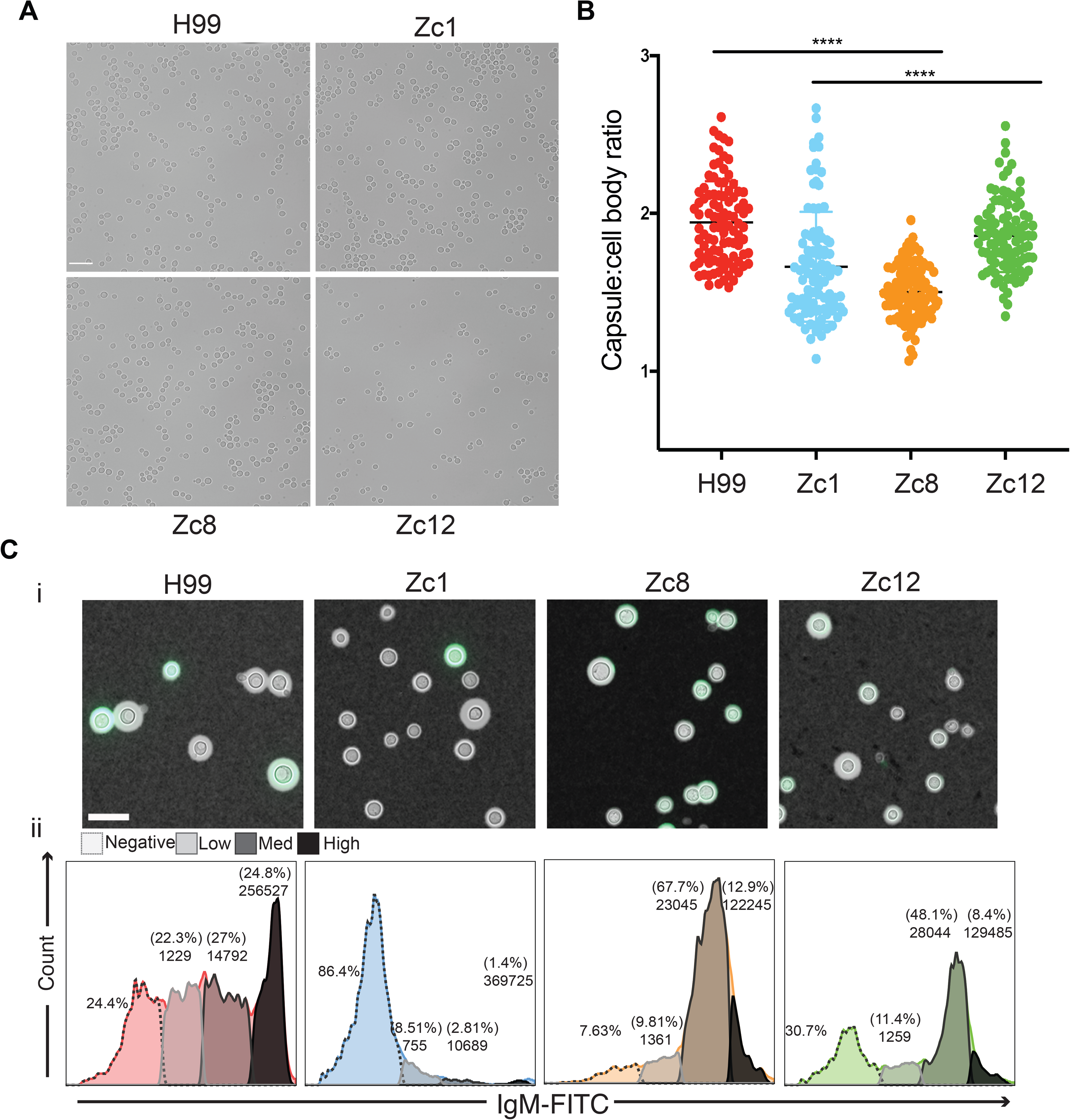
Phenotype characterization of clinical isolates at yeast phase. (A) *C. neoformans* cells from all indicated strains at yeast phase, Scale bar: 20μm. (B) Total capsule and cell body diameters were measured and a ratio reported for each cell. (n>100). ****: p<0.0001.(C) i) Representative images of capsule structure from capsule induction condition stained with both India ink and anti-IgM (Crp127) FITC(Green). ii) Histograms for IgM staining and gates show peaks for different levels of staining with corresponding frequency and MFI reported. Negative: cells failed to bind. Scale bar: 10μm. For all figures, data for a representative biological replicate is presented (n>3).

Simply reducing the cell density leads to titan induction:10% FCS at OD_600_=0.001. After induction, all isolates expressed robust capsule and were reactive to Crp127 IgM. Analysis of capsule to cell body ratio for each of the populations revealed that Zc12 cells display reduced capsule compared to the other isolates (p<0.0001, n>100) (Fig 5B). Across the four isolates, binding by Crp127 remained heterogeneous, but binding patterns changed compared to high density capsule induction (Fig 5A, C), consistent with our observation that Crp127 binding is dynamic throughout titan development [39]. Similar to high-density capsule induction, for H99 and Zc12, more than 50% of the total population was positive for IgM binding, but H99 staining was higher overall, while Zc12 remained more intermediate. For Zc1, which was predominately negative for IgM under high-density capsule conditions, ~40% of cells became positively stained. However, for Zc8 the pattern was reverse: only 5% of Zc8 cells, the largest cells present, were positive for IgM binding, whereas a majority of Zc8 cells were positive for IgM under high density capsule induction (Fig 4C, D). Together, these data suggest that differences in cell density during exposure to inducing conditions (OD_600_ = 0.1 vs 0.001) significantly impact capsule structure, with a high potential for altered capsule expression and organization across genetically related clinical isolates, even in absence of large-scale cell size changes (Zc1, Zc12).

**Fig 5.**
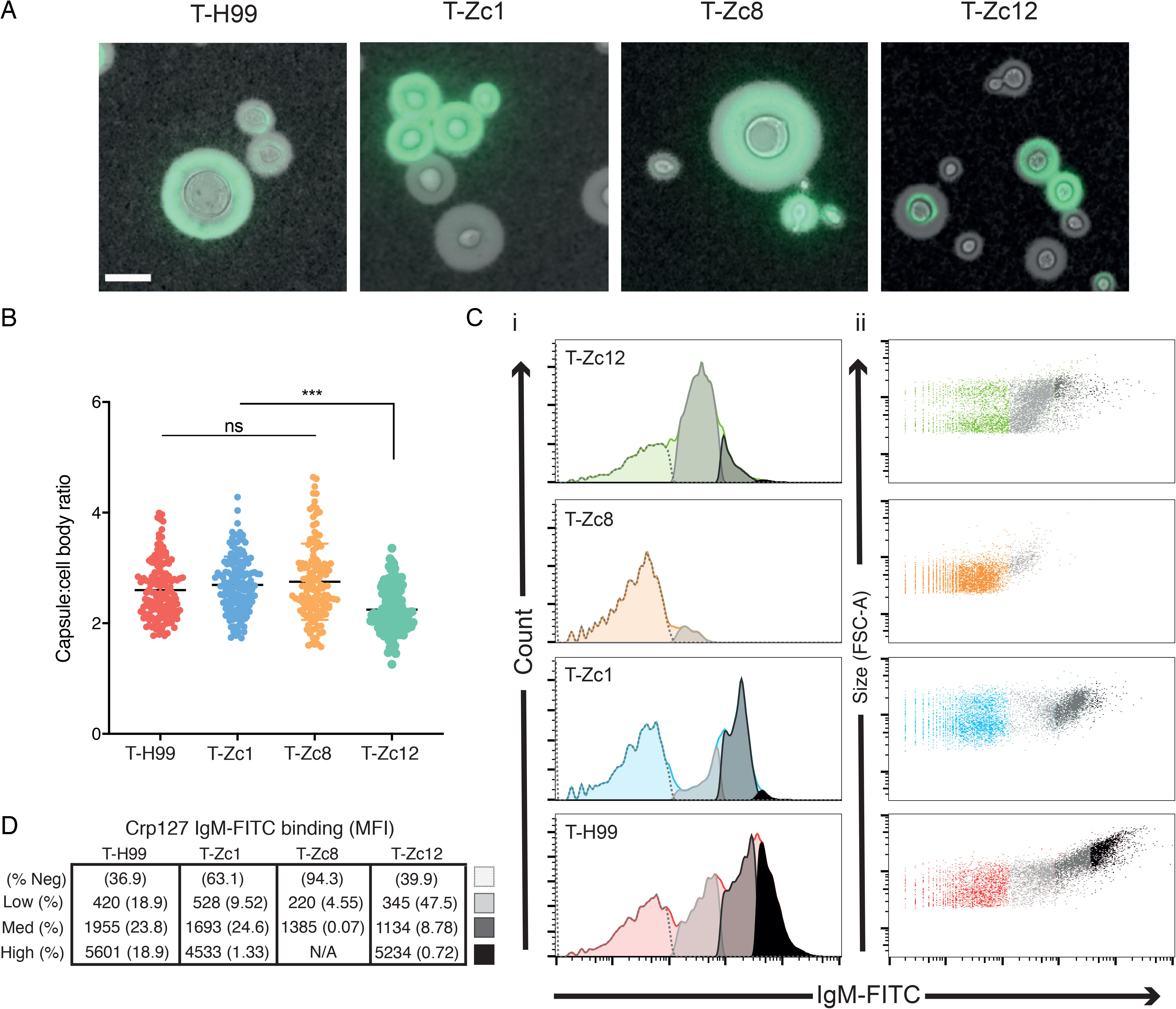
Capsule structure from titan induction. (A) Representative images of capsule structure of titanized cells stained with India ink and anti-IgM (Crp127) FITC(Green); bar scale: 10μm. (B) Total capsule and cell body diameters were measured and a ratio reported for each cell. (n>100). ****: p<0.0001. (C, D) Flow cytometry analysis of IgM staining with frequency and MFI reported. Ci) Gates show peaks for different levels of staining, with (D) corresponding MFI indicated. Cii) Size (FSC-A) vs IgM-FITC intensity, with different populations highlighted as in (Ci). “T” indicates pre-titanized cells. For all figures, data for a representative biological replicate is presented (n>3).

### What are titanides?

Given the lack of correlation between conventional virulence factors and relative virulence in the *G. mellonella* model, we asked whether there was a correlation with the presence of small cells during infection. Specifically, we previously identified a third distinct cell type in induced cultures: oval cells 2-3 μm in diameter with thin cell walls and a clear, well defined nucleus, which we termed titanides [14]. These cells emerge 24 hrs after serum exposure and are the predominant cell type in H99 induced cultures after 3 days (Fig 1A, 6A). We observed titanide cells in Zc8 *in vitro* cultures that shared these features (Fig 1A, 6A) and in *in vivo* lung extracts (Fig S3A), where thick walled “micro” cells could also be observed. Viability analysis of individual cells isolated from *in vitro* cultures by microdissection was 100% within 48 hours of serum induction for both H99 and Zc8 titanides. Titanides are therefore distinct from previously described micro cells (<1 μm) and drop cells (5 μm), which have thick cell walls and, for the latter, disorganized nuclei and low viability [8, 27, 28].

We previously speculated that titanide cells are the daughters of titan mothers. To track their origin, we examined changes in subcellular membrane organization caused by titanization using the ratiometric dye di-4-ANEPPDHQ (Fig 6B)[40]. We observed that the plasma membrane in titan cells is significantly more ordered compared to yeast cells (~5 μm), which had lower lipid packing values (generalized polarization, GP) (Fig 6C). In cells <3 μm, lipids in the plasma membrane were highly ordered, comparable to lipids in titan mother cells, further suggesting these cells are the direct progeny of titan mothers (Fig 6C). Finally, live cell imaging of *in vitro* titan cells revealed that titan daughter cells emerge every 90 minutes and are on average <3.5 μm in diameter (Fig 6D, S4), identifying these as titanide cells.

**Fig 6.**
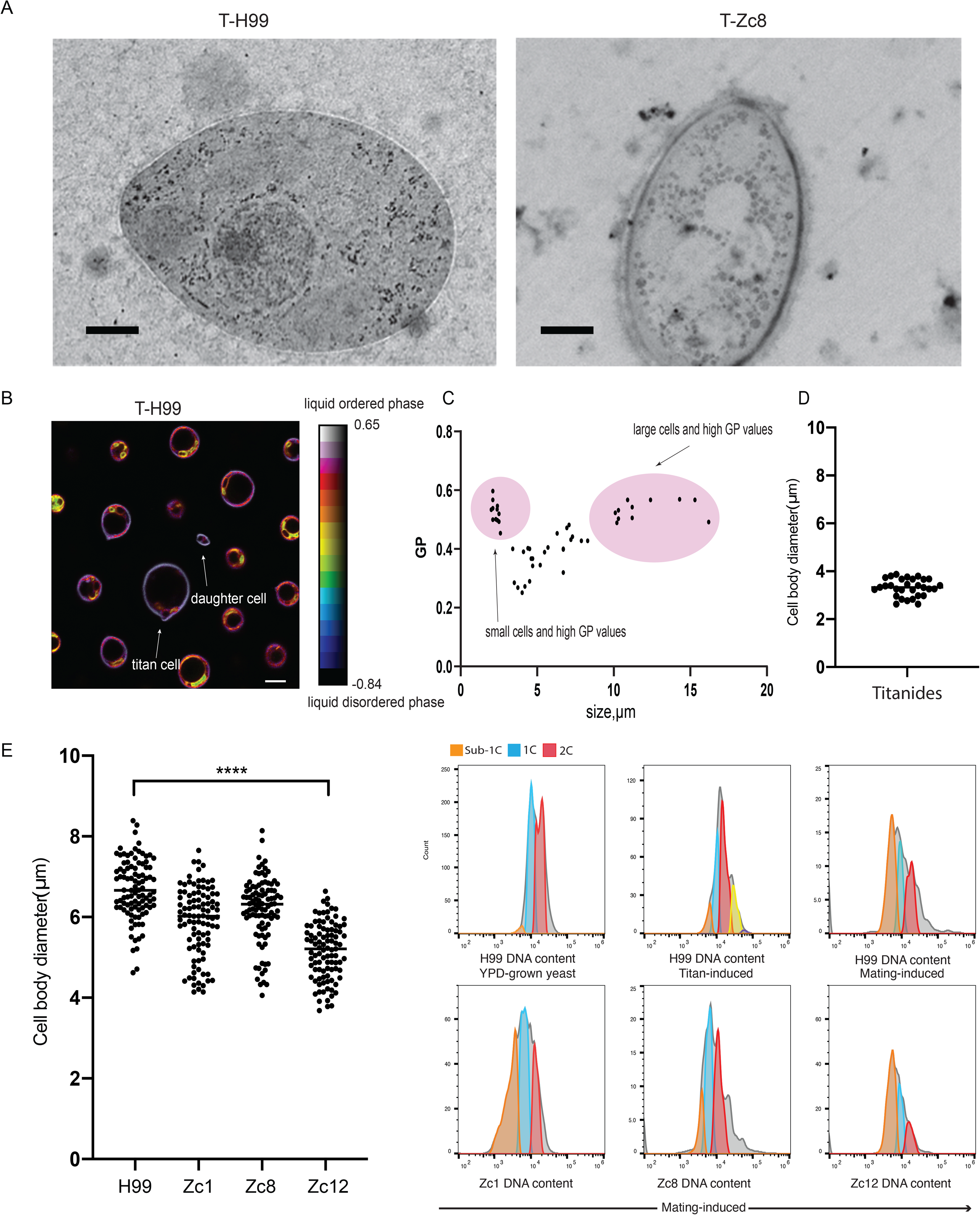
Titanides from both *in vivo* and *vitro*. (A) TEM of titanide cells from H99 and Zc8 from *in vitro* titan induction. Scale bar:500nm (B) Single-plane pseudocolored generalized polarization (GP) images. Color bar designates the range of GP values where red shows high membrane order and blues show low membrane order. Note plasma membrane zones were used to quantify mean GP values. Scale bar: 5μm. (C) A plot representing mean GP values quantified at the plasma membrane region as a function of cell size in H99 cells.(D)Cell body sizes of titan daughters from live imaging of titan cells budding(n=30). (E) Cell body sizes of all indicate strains after capsule induction(n>100). [49] Flow cytometry for DNA content of mating fronts from the indicated isolates crossed with KN99**a**.1C and 2C populations were identified relative to YPD-agar grown yeast cells from matched parent strains and MFI was recorded for sub-1C, 1C, 2C peaks. “T” indicates pre-titanized cells.

Examination of DNA content by flow cytometry for small cell populations revealed that when grown under inducing conditions, all four clinical isolates contained cells with apparent “sub-1C” DNA content, represented as stepwise peaks to the left (Fig 1D). These “sub-1C” populations were not present when cells were grown in YPD (Fig S1 ii). We speculated that these peaks might represent titanide cells, and in fact they are the smallest cells by FSC (Fig S1B). Microscopy to compare cell size in high density capsule inducing conditions vs. low density conditions also showed that these small cells are specifically generated under titan induction (Fig 6E).

Similar “sub-1C” peaks have been observed during flow cytometry-assisted cell cycle analysis of *Saccharomyces cerevisiae* and are representative of G0 or quiescent cells [41, 42]. We speculated that this peak might therefore represent G0 quiescent cells, suggesting that titanides might share similarity to *C. neoformans* spores produced by mating, which are quiescent prior to germination. To test this hypothesis, we first established that quiescent C. neoformans spores exhibit sub-1C DNA content similar to S. cerevisiae quiescent cells. H99, Zc1, Zc8, and Zc12 MATalpha cells were crossed with KN99 MATa cells and incubated for 2 weeks to ensure robust sporulation. Zc1 and Zc12 formed robust hyphae and sporulating basidia after 7 days, comparable with H99. Zc8, a MATalpha isolate, exhibited a profound defect in the formation of hyphae, with sporulating basidia rarely visible even after 3 months (Fig S3B). The mating front, representing hyphae, basidia, spores, and some contaminating yeast, was excised, and filtered through 40 um filters to exclude large particles. The resulting population was fixed, stained and analysed by flow cytometry alongside yeast-phase cells to identify 1C and 2C gates and a “sub-1C” background of <2.5%, set as a threshold for false positives using actively growing YPD cultures. This analysis revealed a “sub-1C” peak to be a major population in robust mating reactions (H99: 37%; Zc1:34%; Zc12:53%) and a minor population (10%) in the Zc8 mating front, in which basidia were rarely observed (Fig 6F, S3B). Direct comparison to titan-induced H99 cultures revealed a similar “sub-1C” peak (8%) suggesting that this population represents quiescent G0 cells specifically generated during titan induction.

### Modelling fungal-phagocyte interaction *in vitro* reveals strain specific phenotypes

To better understand the impact of titan and small cells on disease development, we analyzed the infection process of all four strains following titan induction using an *in vitro* model. We observed significant variation in the percent of macrophages containing at least one internalized *C. neoformans* cell across the four isolates after titan induction, while YNB grown cells were phagocytosed similarly across the four isolates (Fig 7A). H99 showed the lowest macrophage infection rate and was significantly reduced compared to the other 3 strains (Fig 7A) (compared to H99: Zc1, p=0.0063; Zc8, p=0.0131; Zc12, p=0.0164). As shown in Fig 7B, for both H99 and Zc8, engulfed cells were typical cells (titanides and yeasts) rather than titan cells. Furthermore, capsule staining with 18B7 revealed that H99 titan cells physically associated with macrophages but were not engulfed (Fig 7C). We therefore asked whether the reduced uptake of H99 cells overall relative to Zc8 (Fig 7A) is a consequence of the higher proportion of titan cells in this population, similar to in vivo reports. Filtered titan (>10 μm) cells or typical (<10 μm) H99 cells were used to infect macrophages separately.

**Fig 7:**
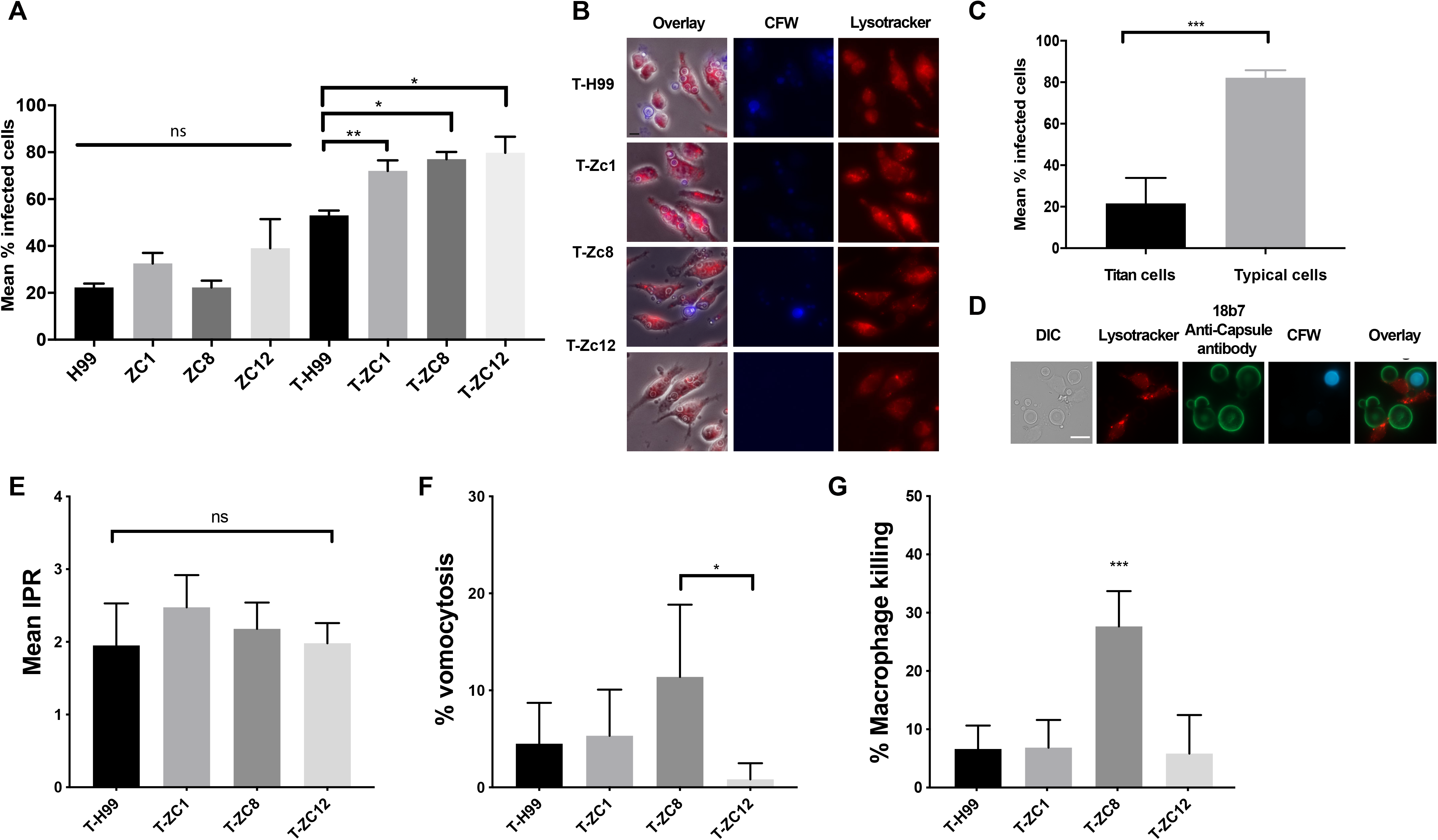
*In vitro* titan cells interaction with host immune cells. (A) Infection rates for the 4 clinical strains, showing the percentage of macrophages infected with at least one internalized *C. neoformans* cell after 2h co-incubation. Data were assessed as not normal by Shapiro-Wilk and analysed by Welch’s ANOVA with Dunnett’s test for multiple comparisons. *p<0.02; ***p=0.0008. (B) Representative images showing macrophages (Red, Lysotracker) infected with cryptococcal cells (Blue, CFW). Scale bar: 10 μm. (C) Macrophages infection rates infected with only titan or typical cells (typical cells <10 um, titan cells > 11 um). Data were assessed for normality by Shapiro-Wilk and analysed by Student’s T-test with Welsh’s correction for unequal variance (determined by F-test). ***p=0.0002. (D) Representative images showing macrophages (Red)infected with H99 yeast and titan cells. Fungi were co-incubated with anti-capsule antibody 18b7 plus anti-mouse IgG-FITC (Green) and calcofluor white (CFW) for chitin (blue). Scale bar: 20μm. (**E-G**) Macrophages and pre-titanized *C. neoformans* were co-incubated for 18 hours and assessed via live imaging for (**E**) IPR, [49] vomocytosis, and (**G**) macrophage lysis. (**E**) IPR: mean intracellular proliferation rate. Data were assessed for normality by Shapiro-Wilk and analysed by One-Way ANOVA. [49] Mf death %: frequency of macrophage death indicated by cell lysis. Data were assessed for normality by Shapiro-Wilk and analysed by One-Way ANOVA. *p=0.05; ***p<0.001. (**G**) Vomo %: frequency of vomocytosis. Data were assessed as not normal by Shapiro-Wilk and analysed by Kruskal-Wallis with Dunn’s correction Data for a representative biological replicate is presented (n>3). “T” indicates pre-titanized cells.

When only titan cells were present, the average macrophage infection was 21.6% ± 5.5 (SEM), while for typical H99 cells the rate was 82.2% ± 1.6 (SEM) (Fig 7D, p=0.0002, n=500), similar to uptake of the total populations for the three Zc strains. Mixed cultures of titanized cells were co-incubated with macrophages in vitro and observed via time-lapse imaging for 18h. Intracellular Proliferation Rate (IPR), frequency of vomocytosis (vomo %), and frequency of macrophage death (Mϕ death %) were measured. No significant difference in IPR was observed for the four strains, indicating that they maintain similar intracellular growth rates (Fig 7E). Zc8 induced the highest frequency of vomocytosis, although it was not statistically significantly different (p=0.05 vs. Zc12) (Fig 7F), but did induce significantly more macrophage death (p<0.001) (Fig 7G; n=400). This suggests that there is no correlation between titanization capacity and vomocytosis or macrophage death. These results are consistent with previous studies *in vivo* showing that titan cells are more resistant to phagocytosis and with the observation that these cells exert a protective effect on the uptake of smaller cells and further validate the overlap between in vitro induction and in vivo induction [5, 12].

### *In vivo* titan inducing conditions match *in vitro* predictions

Overall, *in vitro* phenotyping by microscopy and flow cytometry for cell size and DNA content predicts that: 1) H99 and Zc8 isolates will form titans *in vivo*, with the overall size of Zc8 populations smaller than that of H99, 2) a minority of Zc1 titan cells might be detected, 3) Zc12 will fail to produce detectable titans *in vivo*, and 4) small cells will be apparent for all four isolates and will proliferate to a similar degree within phagocytes. We therefore assessed all four strains in a murine inhalation model of cryptococcosis, the gold standard for determining titan capacity. C57BL/6 female mice were infected intra-nasally with 5×10^5^ cells pre-cultured in sabouroud dextrose medium according to standard infection protocols, and brain and lung fungal burdens were measured after 10 days. Work using H99-derived mutants has demonstrated a role for titan cells in survival and proliferation in the pulmonary environment as early as day 7 [15]. When fungal cells from lung homogenates were analysed microscopically for titanization *in vivo*, similar size distributions were observed as seen in vitro (Fig 8A). Specifically, robust titanization was observed for both H99 and Zc8 (65% and 68% of cells recovered from the lung, respectively, >10 um), with Zc8 population covering a significantly smaller size range than H99 (Kolmogorov-Smirnov test for cumulative distribution, p=0.0001). A minority (8%) of Zc1 cells were >10 um. No Zc12 cells >10 um were observed. Similar to in vitro induced Zc1 and Zc12 cells, the *in vivo* populations as a whole were smaller and more uniform overall. All four clinical strains showed similar fungal burdens in the lungs on day 10 (Fig 8B, p>0.05) as assessed by CFUs, demonstrating similar capacities to proliferate early during infection. Moreover, brain CFUs analysed at day 10 post infection were comparably low across all four isolates, with Zc1 CFUs slightly higher than the others (Fig 8C; p<0.01). Taken together, these results support our predictions that *in vitro* titanization reflects *in vivo* titanization.

**Fig 8.**
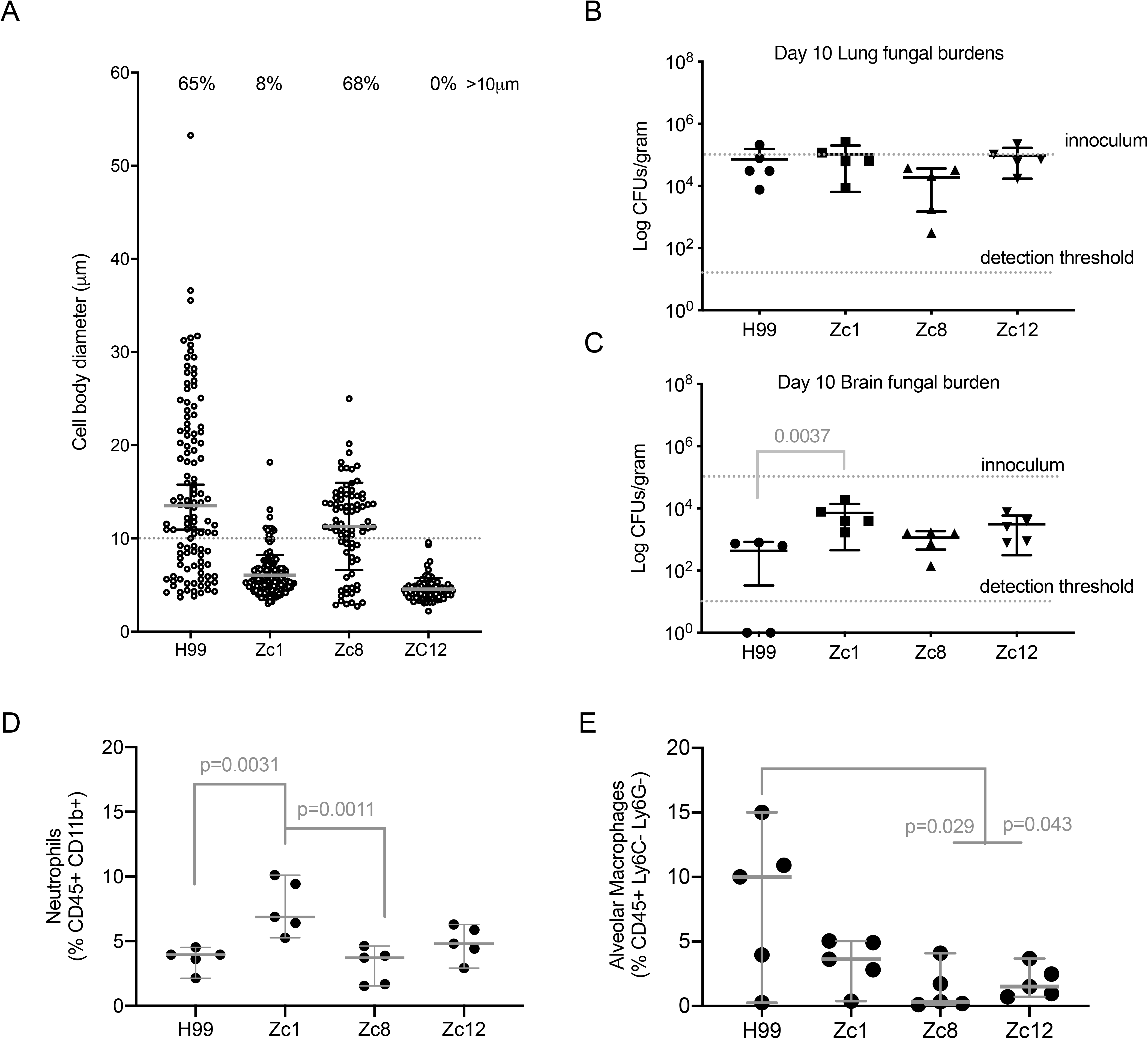
Titanizaiton and disease outcomes on mice model. (A) Cell body sizes of fungal cells collected from murine lungs (n>70 cells for each isolate). (B, C) Infectious burdens for the indicated strains. Murine lungs (B) and brains (C) were assessed for dissemination 10 days post-infection via colony forming units of total homogenates (n=5 per strain). Data are presented as Log CFU/gram. (B) Data were assessed as normal by Shapiro-Wilk and analysed by One-Way ANOVA with Holm-Sidak’s multiple comparisons test. Data are presented as Log CFU/gram. (C) Data were assessed as not normal by Shapiro-Wilk and analysed by Kruskal-Wallis ANOVA with Dunn’s correction for multiple comparisons. (D, E) Immune cell recruitment to lungs for the indicated strains. Lung homogenates were analysed by FACS for the indicated cell populations identified using the indicated markers for (D) Neutrophils (CD45+, CD11b+Ly6G+) or (E) Alveolar Macrophages (CD45+ Ly6C-Ly6G-CD11b-SiglecF+). Data were assessed as normal by Shapiro-Wilk and analysed by One-Way ANOVA with Holm-Sidak’s multiple comparisons test.

### Clinical isolates show different capacities to produce aneuploid colonies

*In vivo* polypoid titan cells were reported to be able generate aneuploid progeny with improved stress resistance compared to typical cells from the same culture [12]. Our *in vitro* model predicts that high titanizing isolates H99 and Zc8 will yield greater aneuploidy *in vivo*. It remains unclear whether ploidy variation in CNS cryptococcosis reflects the degree of ploidy variation observed in the lung. However, because titan daughter cells are smaller on average and readily phagocytosed, we predict that ploidy variation following dissemination to the brain will reflect the degree of ploidy variation observed in the lung. To test this, we determined the impact of titanization on the genetic diversity of offspring and the impact on different sites. We examined the ploidy of individual colonies originating from lung and brain CFUs for all four isolates (Fig 9A, Fig S5, n=30 for lung CFUs, n≥10 for brain CFUs, across 3 mice per isolate). Colonies were assessed within 72 hours of recovery from mice, at the point when a small colony was just visible on YPD agar (~36 generations), a measure of stable aneuploidy in the absence of environmental stress. Fig 9A shows the ploidy of individual colonies relative to the median ploidy of all colonies from that particular lineage. Interestingly, ploidy variation occurred for all isolates, including Zc12. There was no significant difference in relative ploidy between colonies from lung vs. brain isolates for any of the isolates. We observed that ploidy of H99 and Zc8 colonies varied widely, while Zc1 colonies were more uniform (p=0.02 compared to H99). This is consistent with a high degree of titanization driving aneuploidy in H99 and Zc8 infections, and relatively low titanization in the Zc1 isolate. However, we also observed widespread instances of aneuploidy in Zc12 CFUs, despite finding no evidence of true titanization in this strain either *in vitro* or *in vivo*.

**Fig 9:**
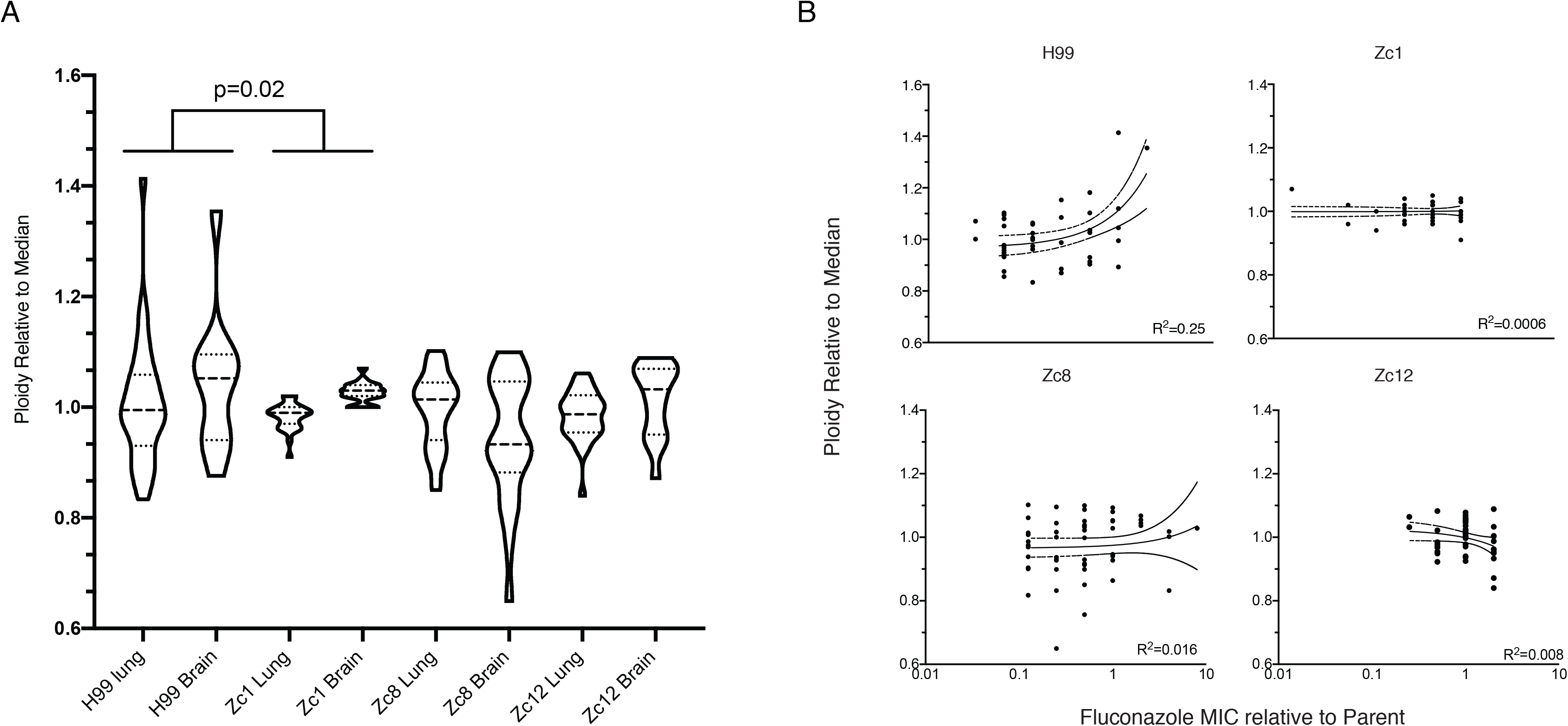
Clinical isolates produce aneuploid daughter cells with altered drug resistance *in vivo*. (A) Degree of heterogeneity in ploidy across the individual colony forming units recovered from host tissue. 1C and 2C populations were identified relative to YPD-grown colonies from matched parent strains and MFI was recorded for 1C and 2C peaks. Data are presented relative to median 1C MFI for all colonies for a given isolate. Data were assessed as not normal by Shapiro-Wilk and analysed by Kruskal-Wallis ANOVA compared to H99 with Dunn’s correction for multiple comparisons. (B) Fluconazole resistance relative to parent for individual colonies aggregated in (A). Data are plotted relative to parent MIC and as a function of colony ploidy as assessed in (A). Data were assessed as not normal by Shapiro-Wilk and analysed by Kruskal-Wallis ANOVA with Dunn’s correction for multiple comparisons.

Aneuploidy is associated with increased resistance to fluconazole, which continues to be used as a frontline antifungal in resource limited settings. Fluconazole resistance in *C. neoformans* is linked to aneuploidy of chromosomes 1, 4, 6, and 10 due to the presence of genes on these chromosomes that directly enable fluconazole resistance, however the contribution of titanization to resistance remains unclear [43]. We therefore assessed the impact of altered ploidy on drug resistance of individual colonies isolated from the lungs and brains of infected mice (Fig 9B). Relative to parent isolates, we observed a wide range in capacity to proliferate in the presence of fluconazole. The distribution of resistance for Zc12-derived colonies was significantly limited compared to those of H99, Zc8, or Zc1 (p≤0.0002). For H99, both fluconazole resistant and sensitive colonies were identified. For Zc8, colonies with increased resistance to fluconazole tended to have increased ploidy, but a colony with reduced overall ploidy and increased drug resistance was also observed. For Zc1, aneuploid colonies were more sensitive to the drug. Within individual lineages there was no correlation between increased or decreased ploidy and degree of fluconazole resistance (R^2^<0.3 in all cases). This is consistent with titanization facilitating ploidy shifts in progeny and with aneuploidy in these isolates being random rather than specific amplification of chromosomes involved in fluconazole resistance occurring prior to drug exposure.

## Discussion

The first clinical and murine model reports of cells consistent with giant/titan cells also reported significant size and morphological heterogeneity. Examination of cells from fungal abscesses revealed a highly heterogeneous population of cells ranging from 2-100 um[9, 10]. Consistent with this, infection of mice with laboratory strains yields a highly heterogeneous population comprised of large, polyploid titans and typical yeast as well as small, thick walled “micro” cells in lung extracts[5, 8, 20, 44]. However, replicating this heterogeneity in vitro presents a challenge to modeling *in vivo* infection. When cultured on rich medium, clinically heterogenous populations become uniform, comprised of only typical yeast[9, 10]. Moreover, this heterogeneity may be site specific within patients: clinical examination of *C. neoformans* isolates taken directly from CSF failed to identify cells that cross the defining 10 μm cutoff [45]. Whether this is because titan cells cannot escape the host lung or because the CSF environment is not supportive of *de novo* titan cell formation remains unclear [34, 46]. Even during co-culture with immune cells that replicate host conditions and elicit titan cells, heterogeneity is not observed [44].

Some degree of heterogeneity is observed under capsule inducing conditions: Fernandes et al. reported that 5 days exposure to DMEM (37°C, 5%CO2) yielded giant and micro cells and rare isolates that showed both giant and micro cells were significantly more virulent[47]. Interestingly, these authors also reported wide variation in size and fungal morphology within and between isolates and across serotypes [13, 47]. Mukaremera et al. also observed that strains that produce small cells were more virulent, using 5 days growth in DMEM+serum (37°C, 5%CO2), conditions in which the high virulence titanizing laboratory isolate KN99a failed to produce titans. However, neither group report observing the wide variation in cell size observed in established titan inducing protocols, nor do they observe small oval cells consistent with titanides[14, 29, 31]. Despite this, these works, together with the findings of Denham et al. reveal a role for small cells in infection progression [19]. While it is established that the presence of titans can affect drug resistance and host immune responses, contributing to poor disease outcome, the role of small cells is less clear [4, 5, 15, 20].

Therefore, a pressing question is the capacity of clinical and environmental isolates to generate population heterogeneity regardless of classic titanization capacity and the effect of this population heterogeneity on disease outcome.

### Investigating population heterogeneity *in vitro*

Here, we expand upon the *in vivo* definition of titan cells to encompass the wide phenotypic diversity observed during *in vitro* induction of three clinical isolates from the ACTA Lusaka Trial in Lusaka, Zambia [26, 33]. Our data add to existing knowledge by specifically studying clinical isolate titanization using an established in vitro titan induction protocol. After in vitro titanization, *C. neoformans* populations are highly heterogeneous with multiple sub-populations: titan (>10um), typical/small (>4 μm), and titanide (2-4 μm) cells, as well as micro (1 μm) cells [5, 8–10, 14, 44]. Building on our previous findings, we used 4 clinical strains from the same genetic clade (VNI) which we previously identified as having different capacities to produce titan cells [14]. These four strains are closely related based on genome analysis, however evolved variation still exists between them.

All 4 strains demonstrated significant population heterogeneity after exposure to inducing conditions, regardless of titan status. Relative to Zc8, H99 is strongly titanizing *in vitro* when the *in vivo* definition (> 10 μm) is applied, but Zc8-induced cells exhibit overall population structure consistent with titanization, including polyploidy (>4C), a relative increase in cell size, and overall changes in PAMP organization. By combining microscopy and flow cytometry, we propose a model in which predictive outcomes can be made about *in vivo* strain “titan-signature” status. We additionally note that researchers can readily identify titan cells through visual inspection of liquid cultures in tissue culture plates using bright-field microscopy, with titans being large cells with a single vacuole and an intensely dark cell wall apparent in H99, Zc8, and Zc1 cultures, but DNA content analysis offers important information about overall population structure (Fig 1A).

### Variation in cell size and morphology and impact on host cell interaction

Population heterogeneity appears to be a general feature of titanizing isolates. Here we observed that both titanizing strains H99 and Zc8 are highly heterogeneous after induction, with three sub-populations that maintain distinct cell sizes and DNA contents. Moreover, titan cells from both strains exhibit changes in PAMP exposure and capsule size and composition, features important for the virulence of *C. neoformans*. Surprisingly, non-titanizing induced strains Zc1 and Zc12 also show population heterogeneity as measured by DNA content and cell wall structure, despite a lack of true titan cells. Consistent with previous findings, we directly showed that the presence of titan cells leads to decreased phagocytosis rates of small cells [5, 12]. Denham et al. showed that at later infection stages, in both lungs and brains the predominant population is the smaller cryptococcal cells around 10μm in total diameter[19]. This matches our *in vitro* observation that in later titan-induced cultures, the proportion of small cells increases, consistent with the increased replication rate of titan mothers and the impact of cell culture density on the generation of new titan mothers[14, 29, 31].

We additionally observed changes in capsule and IgM binding upon titan induction relative to other capsule inducing protocols. IgM mediates complement-dependent phagocytosis and has been shown to specifically inhibit the yeast-to-titan transition [48, 49].Interestingly, IgM has been shown to be particularly important in the clearance of small particles (2-5μm) [50]. Our data here suggest differences in epitope exposure on small vs. large cells may exert a protective effect by reducing overall phagocytosis. Future work will examine the impact of titan induction on complement-dependent, opsonin-independent uptake and will address the relative pathogenic potential of titanide vs yeast vs spores in murine models of infection.

### How do titanides contribute to disease progression?

There is growing evidence of a high degree of genome plasticity during *C. neoformans in vivo* growth [51–53]. Semighini *et al*. previously demonstrated, using comparative genome hybridization arrays in H99-derived strains, that passage through mice in the absence of drug stress results in large scale genome alterations, including aneuploidy, insertions, and deletions [51]. Gusa *et al*. recently demonstrated that host temperature is sufficient to induce transposon-mediated rearrangements in the *C. neoformans* genome [52].However, titanization adds to and enhances this variation: titan cells produce aneuploid daughter cells that promote *in vitro* stress resistance relative to yeast in the same population [12]. We add to these previous reports by showing that high titanizing H99 and Zc8, low-titanizing Zc1 and non-titanizing Zc12 isolates all yield aneuploid daughters to some degree when exposed to the host microenvironment. We reveal that titanization can drive the emergence of aneuploid offspring in the host by comparing the ploidy distribution of cells recovered from infected mice. We observed large scale changes in genome content relative to parent isolates, and this weakly correlates with degree of titanization (H99>Zc8>Zc1>Zc12). We hypothesize that, in addition to strongly inducing titanization, during infection, the host microenvironment itself is sufficient to induce aneuploidy and exerts a selective pressure, as has been observed for *C. albicans* [54]. In line with this, we previously reported that daughter cell colonies derived from *in vitro*-induced titan cells are mostly diploid, in conflict with reports from Gerstein *et al*. that colonies derived from *in vivo*-derived titan cells are haploid or aneuploid [12, 14].

This suggests that daughter cells derived from the two conditions may differ in their tolerance to aneuploidy or capacity for genome maintenance. Fluconazole exposure, oxidative stress, and high temperature drive *in vivo C. albicans* aneuploidy, including whole ploidy reduction (diploidy to haploidy), individual chromosome aneuploidy (Chr 5), and segmental aneuploidy, and the same is likely true for *C. neoformans* even in the absence of titanization [54–57].

Future work will more fully probe the molecular mechanisms differentiating these clinical isolates and their impact on virulence.

### What is the environmental significance of “sub-1C” cells?

Ashton et al. recently suggested that the existence of robust quiescent cells such as spores might contribute to the broad ecological distribution of specific sub-clades, however the rarity of MATa *C. neoformans* isolates in the environment suggests an alternate origin for these cells[58, 59]. Here, we demonstrate that titanides observed in induced cultures originate from titan mothers and that these cells exist in a in G0-like state similar to spores generating by mating. Titanide cells have unique features including oval shape, smaller size and altered DNA content compared with typical yeast cells. The ability to induce titan cells in conditions likely to be encountered by Cryptococcus in the environment make it plausible that titan cells may form in the environment and are not just limited to patient lungs. In addition, we also we detected “sub-1C” populations from the Zc12 isolate in titan-inducing conditions, suggesting that the capacity to produce quiescent cells is not simply a consequence of high ploidy mothers. This raises the hypothesis that “quiescent” cells proposed to drive population expansion in environmental isolates may be generated via regulatory pathways driving titanization [58].

### Impact of population heterogeneity on drug resistance

Gerstein *et al*. previously demonstrated that *in vivo*-derived titans provide daughters with a survival advantage compared to *in vivo*-derived typical cells[12]. Therefore, we predict the presence of titanides also contributes to disease progression by affecting drug resistance and stress adaption. Titan-derived colonies are resistant to fluconazole via aneuploidy of chromosomes 1, 4, 10, 11, and/or 12 when propagated in the presence of fluconazole, suggesting that titanization drives fluconazole resistance *in vivo* [12, 43]. Our data support these findings, with overall increases in population diversity in titanizing strains compared to non-titanizing strains following passage through mice (Fig 7A). However, we observed no specific correlation between increased ploidy and increased fluconazole resistance (Fig 7B).

This is consistent with *in vivo*-induced aneuploidy being random with regard to particular chromosomes, rather than a specific increase of chromosomes involved in fluconazole resistance. Finally, induced but not un-induced cells from all isolates showed high but similar virulence when using the galleria model and we found no differences in capacity to control early lung fungal burdens during a mouse model of cryptococcosis. Our data therefore demonstrate that in response to the host environment signals both non-titanizing and titanizing isolates generate heterogeneous populations which will have implications for drug resistance and host cell interactions. Future studies will investigate their impact on long term immune responses including investigation of T cell polarization mediated by the different clinical isolates.

## Acknowledgements

We thank the doctors, patients, and their families involved in the ATCA Lusaka trial, without whom this research would not have been possible. We thank the staff of the University of Birmingham BMSU and the University of Aberdeen Medical Research Facility for their assistance. 18B7 anticapsule antibody was a generous gift from Arturo Casadevall. Crp127 IgM anticapsule antibody was a generous gift from Robin May. We are grateful to Debbie Wilkinson and Lucinda Wight in the University of Aberdeen Microscopy and Histology Core Facility for their expert help with TEM experiments.

## Materials and methods

### Strains, media and growth conditions

All *C. neoformans* strains used in this study are listed in Table 1.

| Strains | lineages | Clades | Source |
| --- | --- | --- | --- |
| H99 | VNI | Clinical isolate | North Carolina |
| Zc1 | VNI | Clinical isolate | Zambia |
| Zc8 | VNI | Clinical isolate | Zambia |
| Zc12 | VNI | Clinical isolate | Zambia |
| CNV111 | VN1 | H99-derived | [60] |

Strains were cultured routinely on YPD (1% yeast extract, 2% bacto-peptone, 2% glucose, 2% bacto-agar) plates stored at 4°C. For mating reactions, cells were mixed from YPD plates with KN99a directly on Murashige Skoog agar (M5519 Sigma-Aldrich) and incubated in the dark at room temperature for at least 2 weeks. Titan cell *in vitro* induction was performed as described in [14]. Briefly, cells were pre-cultured at 37°C, 150rpm, in 10ml YNB without amino acids (Sigma-Aldrich) in which 2% of glucose was added according to manufacturer’s protocol. After 24 hrs, cells were normalized to OD_600_=0.001 in 1XPBS (phosphate-buffered saline) with 10% of Heat inactivated Fetal Bovine Serum (HI-FBS) (Sigma-Aldrich F9665) and incubated at 37°C, 5%CO_2_ for 24 hrs unless otherwise indicated.

### *Galleria mellonella* infection model

*G. mellonella* were purchased from LiveFoods, Ltd (UK) and used within 3 days of arrival. Worms were maintained at 16° C until use, and only worms lacking melanization were included in the experiment. Worms were allocated randomly to each treatment group (10 per group), with the average weight of worms across treatment groups maintained within 0.002 g. Cryptococcus strains were incubated overnight in YPD (5% yeast extract, 10% bactopeptone, 2% D-glucose) at 150 rpm, 30 ° C, or YNB (and then exposed to 10% serum in PBS at OD_600_, 5%CO_2_, 37 ° C for 16 hrs. Cells were washed in sterile PBS, and then normalized to 5× 10^6^/ml. Strains were randomly allocated to the different groups. Worms were injected into the back left pro-leg with 20 ul innoculum. Worms were maintained in the dark at 30 ° C and observed for 10 days for survival, with blinding for the identities of the different treatment groups. Survival was analyzed by Kaplan-Meyer survival curve and significance was assessed by Log rank (Mantel-Cox) test.

### Murine infection model

Female C57BL/6 mice were used at 8-10 weeks of age and were maintained in individually-ventilated cages under specific pathogen-free conditions at the Biomedical Research facility at the University of Birmingham (Birmingham, UK). Animals were provided with food and water *ad libitum*. All experimentation conformed to the terms and conditions of United Kingdom Home Office license for research on animals (PPL/PBE275C33) and the University of Birmingham ethical committee.

### Cryptococcosis Infection Model

Yeast was grown in SabDex broth (Sigma-Aldrich), grown at 30° C with shaking for 18-24 hours. Yeast cells were washed in PBS, counted, and delivered to mice intranasally while under isofluorane anaesthesia. Animals were infected with 2×10^5^ CFU. For analysis of lung and brain fungal burdens, animals were euthanized and organs weighed, homogenized in PBS, and serially diluted before plating onto YPD agar supplemented with Penicillin/Streptomycin (Invitrogen). Colonies were counted after incubation at 37° C for 48 hours.

### Preparation of Lungs for FACS

Lungs were removed aseptically from euthanized animals, finely minced with a scalpel, and digested in digest buffer (RPMI supplemented with 10% FBS, 1% penicillin/streptomycin [Invitrogen], 1mg/mL collagenase D [Sigma], 1mg/mL dispase [Sigma], 40 μg/mL DNAse [Roche]) at 37° C for 30 minutes. Digested lungs were smashed through a 70μM filter, washed and red blood cells lysed on ice using BD PharmLyse solution. Lung-isolated cells were washed in PBS/2mM EDTA, and then stained with fluorophore-conjugated antibodies in the presence of anti-CD16/32 and 0.5% BSA for 30 minutes on ice. Samples were washed in PBS/0.5% BSA/0.01% sodium azide and acquired using the BD Fortessa instrument equipped with BD FACS Diva software (BD Biosciences). FlowJo (TreeStar) was used for the final analysis. Anti-mouse antibodies used in this study were: CD45 (30-F11), Ly6G (1A8), Ly6C (HK1.4), SiglecF (S17007L), all from Biolegend, and CD11b (M1/70), F4/80 (REA126), MHC Class II (REA813), all from Miltenyi Biotec.

### Cell wall staining

Cells from 24 hrs induced titan cultures were collected, washed twice with 1xPBS and resuspended in 500ul McIIvaine buffer. Cells were stained with EosinY (5μg/ml) in dark for 10mins and repeated for washes with McIIvaine buffer, then resuspended in 500ul 1XPBS. CFW (10 ug/ml) and ConA (50 ug/ml) were added into samples and maintained in the dark for 10 mins. After washing twice with 1XPBS, cells were imaged with a Zeiss Axio Observer and analysed for fluorescent intensities with Attune NxT Flow cytometer.

### J774.1 cell culture

For most *in vitro* infection assays, we used the J774.1 murine macrophage cell line. Cells were routinely passaged in Dulbecco’s modifies Eagle medium (DMEM+) culture media with serum (DMEM, low glucose, from Sigma-Aldrich; 10% HI-FBS; 1% penicillin-streptomycin solution from Sigma-Aldrich; 1% 200mM L-glutamine from Sigma-Aldrich). All assays were performed with cells in passages 4 to 10.

### *In vitro* phagocytosis assays

The intracellular proliferation assay was performed as previously described [61]. 24h before infection, 1×10^5^macrophages J774 were seeded into 12-well plastic plates [62] in 1 ml DMEM culture media with serum (DMEM+) and activated with 10 U/ml IFNγ. Before infection, *C. neoformans* cells were collected from titanization culture and washed three times with sterile 1XPBS. After normalising to 1X10^6^ cells/ml in 300 μl DMEM+, cells were opsonized with 10μg/ml anticapsular 18B7 antibody by incubating at 37°C for 30min [63]. Then opsonized *C. neoformans* cells were added into activated J774.1 macrophages to give a multiplicity of infection (MOI) ratio of 10:1. After 2 hrs (time point 0), infected wells were washed at least three time with pre-warm 1XPBS until all un-engulfed *C. neoformans* cells were removed from the wells. Human monocytes were infected with opsonized cells for 4h without IFNγ activation, and at this point all supernatants were collected for cytokine qualification. 4h post-infection, monocytes were divided into two groups: with or without IFNγ (10 U/ml) activation and then infection continued for another 20h. Total 24h later, all supernatants were collected again with LPS (100ng/ml) as a positive control added at the same time as fungal cells and uninfected as negative control.

### Macrophage infection rates

At time point 0, attached *C. neoformans* cells and macrophages were stained with Calcoflour white (CFW, 10 μg/ml) and Lyso-Tracker Red (ThermoFisher, 1 μM/ml) in dark for 10 min and then fixed with 1 ml 4% paraformaldehyde (PFA) in 1XPBS at 37°C for 5min. After washing with 1XPBS, at least 5 frames from each single infected well were randomly selected and imaged with Nikon Ti-E microscope, capturing 500 cells. All images were analysed with FIJI and experiments were repeated 3 times for biological replicates.

### Filtered Titans phagocytosis assays

After 24h of *in vitro* induction, H99 titanised total population was passed through an 11μm filter to separate titan (>11μm) and typical cells (<11μm). Then both populations were washed, normalized and opsonize independently before adding to infection assays. Activation and infection of macrophages with opsonized *C. neoformans* cells followed the same protocol as described above.

### Intracellular proliferation and vomocytosis assays

At time point 0 of phagocytosis assays, extracellular cells were removed by washing with warm fresh DMEM+. All samples were maintained at 37°C, 5%CO_2_ in the Nikon Ti-E microscope chamber and imaged for 18h via time-lapse. Images were taken every 15 min and compiled into single movie files for analysis using NIS elements or FIJI software. Movies then were scored visually for intracellular proliferation rate, macrophage death and vomocytosis percentage. Vomocytosis scoring followed the guidelines described in Andrew et al., 2017 [64].

### Flow cytometry for cell ploidy

Cells were fixed and stained according to the protocol of [14]. Cells were fixed with 50% methanol for 15 mins, washed 3 times with 1XPBS and stained with 3 ug/ml DAPI. Cells were analysed for DNA content using an Attune NxT Flow cytometer with violet laser. All flow cytometry data were analysed with FlowJo. Doublets and clumps were excluded using recommended gating system of FSC-H vs FSC-A, which was determined to be more parsimonious than SSC-H vs SSC-W followed by FSC-H vs. FSC-W. Autofluorescence in the DAPI channel was excluded by comparison to unstained cells. Gates for 1C and 2C were established with haploid controls cultured in YPD liquid media as described in the text.

### Cell sizes measurement

Cell diameter was measured using FIJI, with frames randomly selected, all cells in a given frame analyzed, and at least 5 images acquired per sample for each of two independent runs. Total cell number of each samples was >200. All statistical analyses were performed using Graphpad Prism.

### Measurements of membrane lipid order *in vivo*

Cells from 24 hrs induced titan cultures were collected, washed twice with 1xPBS. Following resuspension in fresh medium containing 5 μM di-4-ANEPPDHQ, cells were incubated for 30min at 30°C. Cells were then transferred into a glass-bottomed microscope dish. Imaging was performed on a Zeiss LSM 780 confocal microscope equipped with a 32 element GaAsP Quasar detector. A 488 nm laser was selected for fluorescence excitation of di-4-ANEPPDHQ. The detection windows were set to 510–580 nm and 620–750 nm. The images were analyzed using a plug-in compatible with FIJI.

## Supplemental Figures

**Fig S1. Supplemental information for DNA content analysis used Fig 3**. (A) i) Histograms of DAPI staining intensity for YPD grown cells, and 1C and 2C peaks are indicated based on the MFI. ii)FSC-A vs DNA content scatter plots show the correlation of size increase with DNA content. (B) Single cells were identified by FSC-A vs FSC-H to remove doublets (n>8000 after gating), then were used for DNA content analysis (Left panel). FSC-A vs DNA content scatter plots show the correlation of size increase with DNA content (Right panel). “T” indicates pre-titanized cells.

**Fig S2.** (B) PAMPs staining flow cytometry analysis of induced cells from Fig 8 with large cells sub-population highlighted in grey and MFI indicating exposure of Chitin (CFW, right); Mannan (ConA, center); and Chitosan (EosinY, left). “T” indicates pre-titanized cells.

**Fig S3.** (A) TEM of Zc8 titanides recovered from mice lungs, scale bar: 500nm. (B) Mating plates from the indicated isolates crossed with KN99 MATa.

**Fig S4.** Live imaging of CNV111, expressing Nde1-GFP and Cse4-mCherry in an H99 background. Cells were incubated overnight in YNB, and then incubated in 10% FCS, 5% CO_2_, 37°C at OD_600_=0.001 for 6 hours. Cells were immobilized in glass-bottom chamber wells using 1:200 dilution of Crp127 and imaged using a Zeiss Axio-imager at 63x, 36°C, 5%CO_2_. Images were acquired every 10 minutes. The movie is representative of >10 biological repeats.

**Fig S5**: DNA content of individual colony forming units recovered from host tissue, L-M1: Lung of mice 1, B-M1: Brain of mice 1, dotted lines indicate median 1C MFI. n=24, except H99 samples from brains(n=21) because of limitation of colonies recovered.

## Notes

Funding XZ is supported by a studentship from the Darwin Trust. HZ is supported by an NC3Rs PhD studentship (NC/R001472/1). PSC was funded by a BBSRC MIBTP PhD studentship. RAD and SHM are supported by the Academy of Medical Sciences (SBF004_1008) and the Medical Research Council (MR S024611_1). MM is funded by the University of Birmingham School of Biosciences. AC and DMM were supported by the NC3Rs (NC/N002482/1). IMD is funded by the Wellcome Trust (102705 and 097377) and the MRC Centre for Medical Mycology (MR/N006364/1). ERB is supported by a Sir Henry Dale Fellowship jointly funded by the Wellcome Trust and the Royal Society (211241/Z/18/Z).

### Competing Interest Statement

The authors have declared no competing interest.

### Summary of Updates

Substantial revision of the text and includes additional experimental data based on peer-review feedback. We thank the anonymous reviewers who made this possible.

