## Supplementary figures and images for "Host environmental conditions induce small fungal cell size and alter population heterogeneity in *Cryptococcus neoformans*"

### Fig S1

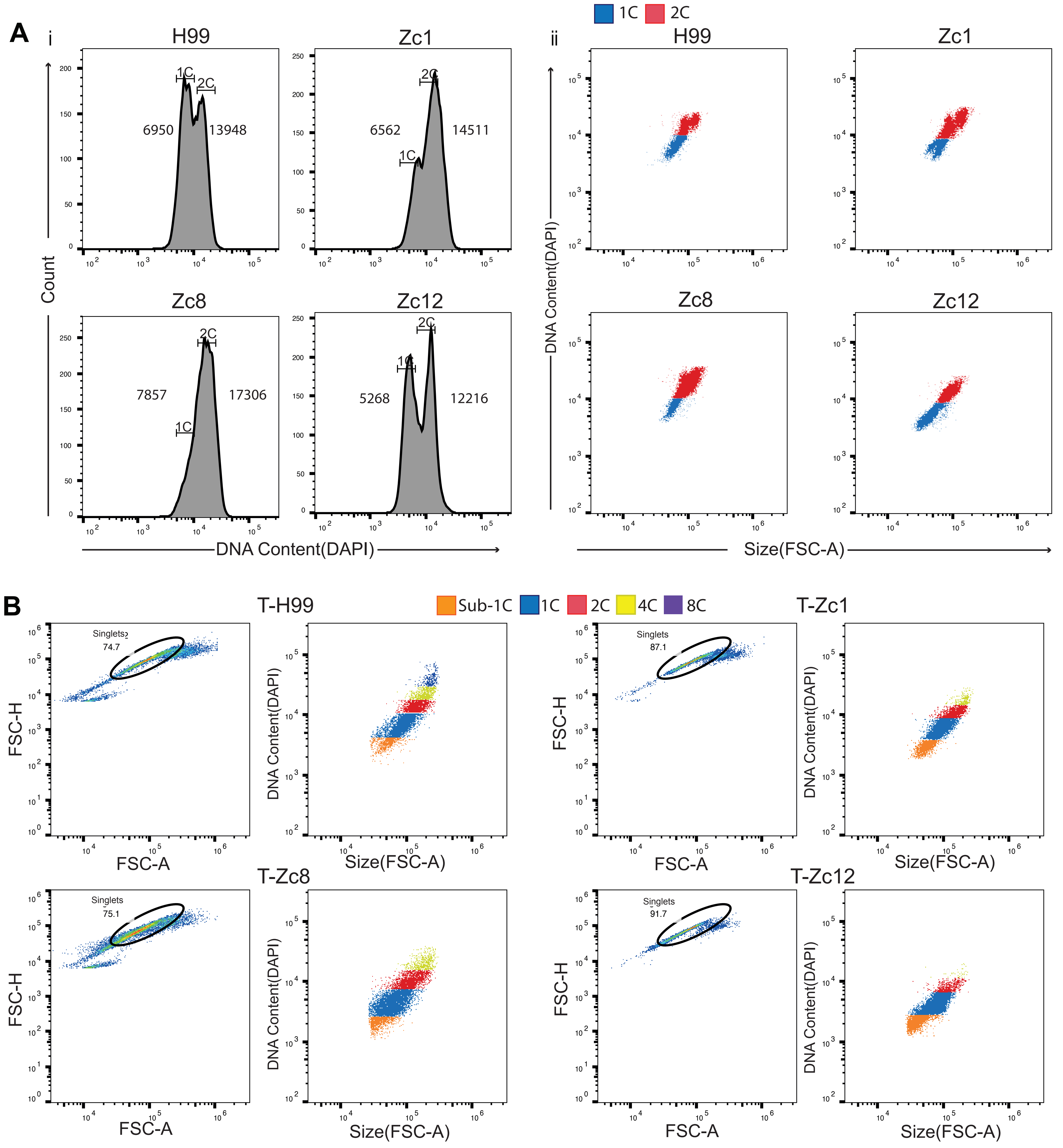

### Fig S2

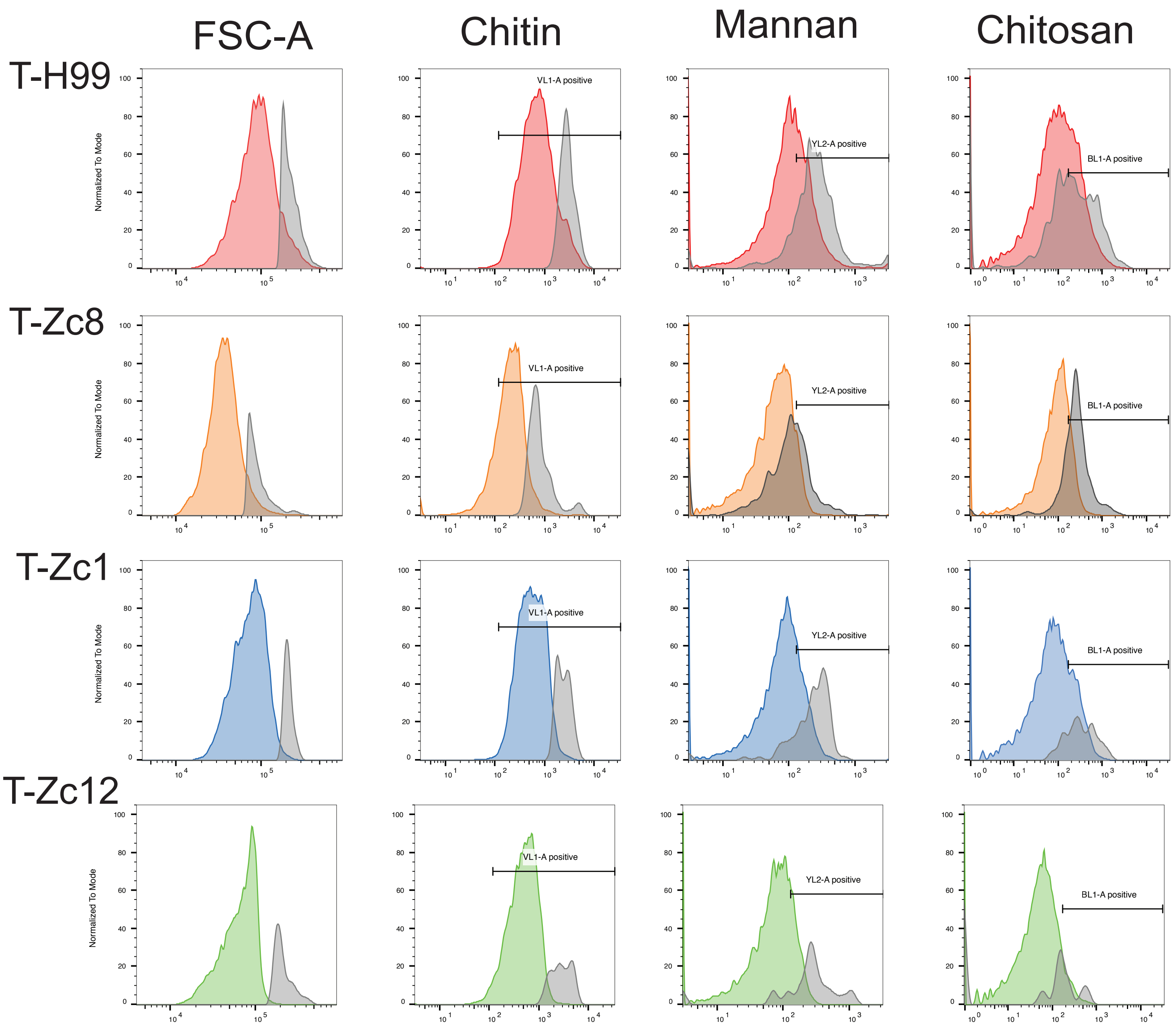

### Fig S3

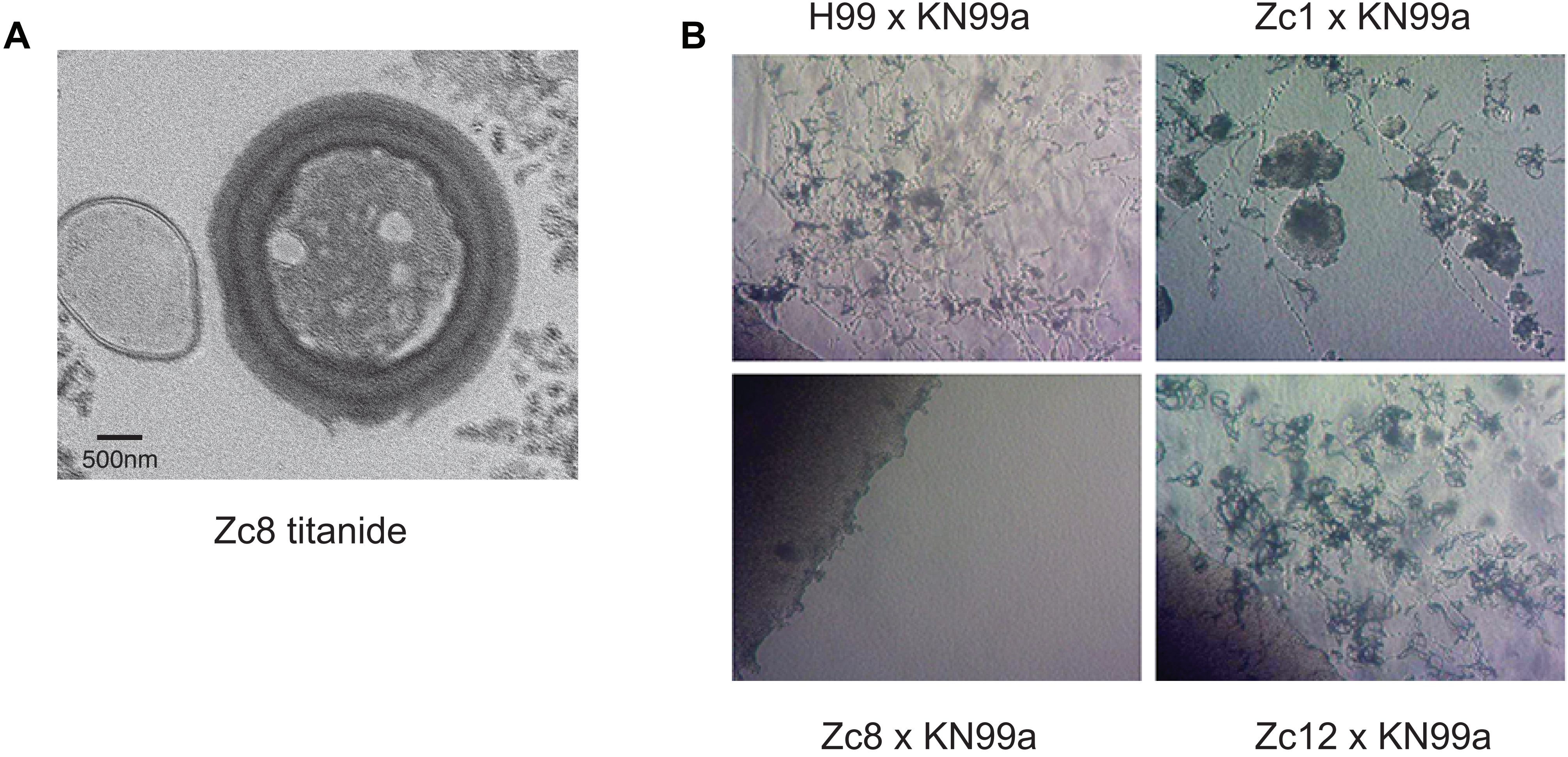

### Fig S5

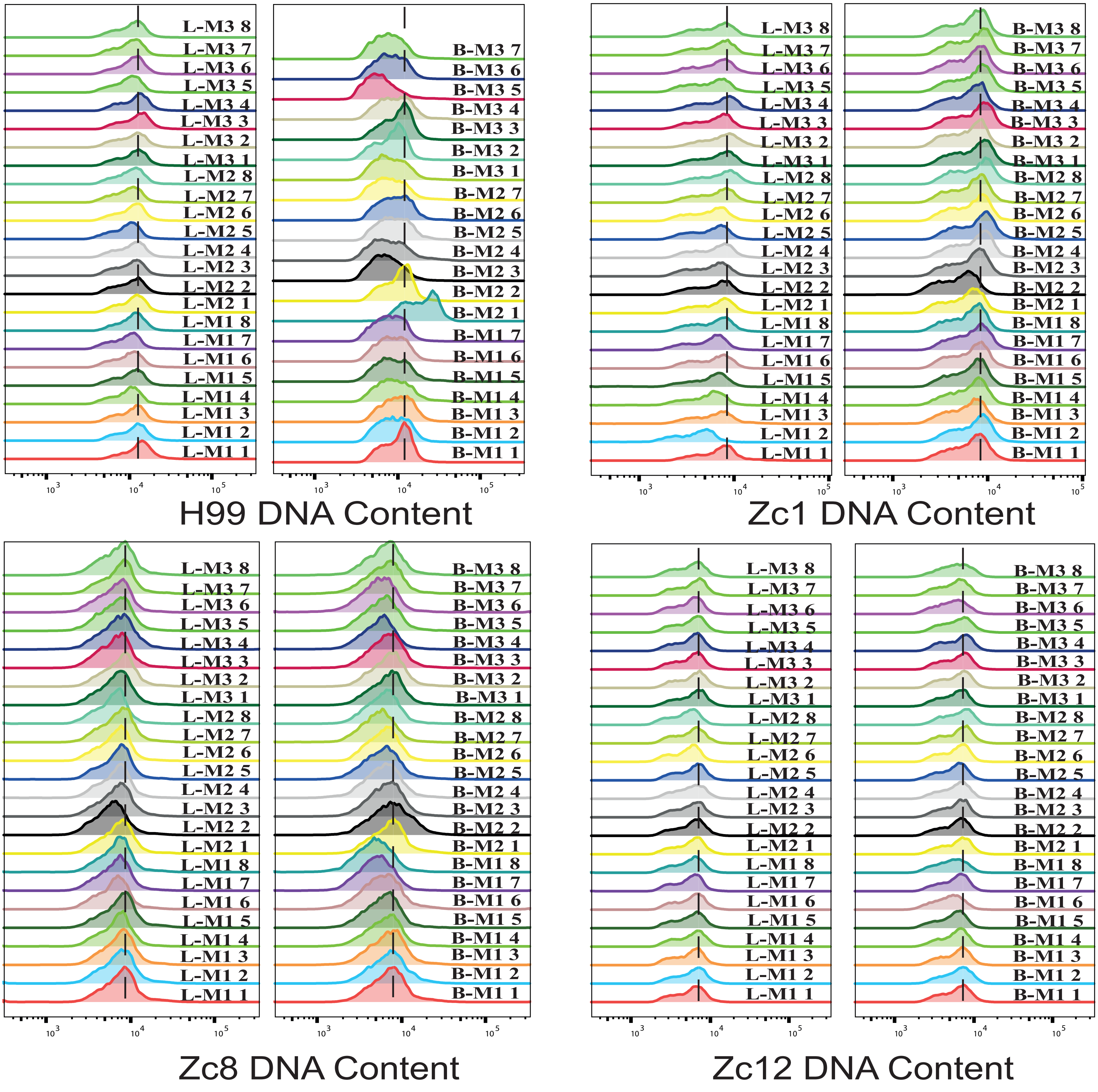
